# Restriction enzymes use a 24 dimensional coding space to recognize 6 base long DNA sequences

**DOI:** 10.1101/538025

**Authors:** Thomas D. Schneider, Vishnu Jejjala

## Abstract

Restriction enzymes recognize and bind to specific sequences on invading bacteriophage DNA. Like a key in a lock, these proteins require many contacts to specify the correct DNA sequence. Using information theory we develop an equation that defines the number of independent contacts, which is the dimensionality of the binding. We show that EcoRI, which binds to the sequence GAATTC, functions in 24 dimensions. Information theory represents messages as spheres in high dimensional spaces. Better sphere packing leads to better communications systems. The densest known packing of hyperspheres occurs on the Leech lattice in 24 dimensions. We suggest that the single protein EcoRI molecule employs a Leech lattice in its operation. Optimizing density of sphere packing explains why 6 base restriction enzymes are so common.

## 1 Introduction

Restriction enzymes provide a defense mechanism in procaryotes against foreign DNA injected by bacteriophages [1,2]. These proteins bind to specific sequences on DNA and cleave the DNA, rendering it susceptible to attack by exonucleases [3, 4] and preventing viral replication. The bacterial genome is protected by modification enzymes that methylate the same pattern that the restriction enzymes cut. Though the sequences bound by restriction enzymes usually consist of only 4 or 6 base pairs, even a single base change of the GAATTC EcoRI binding site decreases EcoRI binding by at least 1000 fold [5]. How can restriction enzymes have such precise recognition? Why do we find the majority of restriction enzymes have exactly 4 or 6 base pair long recognition sequences?

We begin with a brief overview of how concepts from information and coding theory can be used to answer these questions. We address this paper to both biologists and information/coding theorists. Therefore some of this material may be familiar to one audience and foreign to the other.

In this paragraph, we state our main result. We model the binding process of restriction enzymes as a selection between distinct states. These states can be represented as spheres packed together in a high dimensional ‘coding’ space, and the equations of information theory along with empirical data for EcoRI binding allow us to determine the dimensionality of the space. Surprisingly, operating at the maximum biological efficiency, the 6-base cutting enzyme EcoRI works in a 24 dimensional space, and so is likely to use the best sphere packing known, the famous 24 dimensional Leech lattice [6]. Using the Leech lattice would allow restriction enzymes to minimize their errors and so cut precisely. The 4-base cutters should work in a 16 dimensional space, and in that dimension there are also good sphere packings. Apparently, restriction enzymes have evolved over a high dimensional landscape to take advantage of the best sphere packings.

To model recognition of EcoRI binding to DNA, we distinguish two different energy flows in time. The first is the energy dissipated during the binding or ‘operation’ of the molecules *P* (power) [7]. To describe *P* for EcoRI we consider two states, and in both states the protein is associated with the DNA. The dissipation of *P* proceeds from a high energy ‘before’ state in which the EcoRI molecule is somewhere on the DNA but not bound specifically. Then, after a Brownian motion search [8,9] when EcoRI encounters a binding site it may begin to form specific bonds [10,11]. As these bonds form, energy is dissipated to the surrounding water until EcoRI has formed all its bonds to the DNA. We call this latter low energy state the ‘after’ state. The energetic difference between these two states is the specific binding energy, 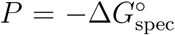 with units of joules per binding [12]. The time of this dissipation may vary, but the total energy dissipated is constant between the two states, as indicated by the measurability of 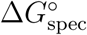 for the operation [13]. The second important energy flow involved in EcoRI binding is the thermal noise that passes through the molecule *during the binding operation*. This noise, *N*, interferes with bond formation. These concepts are parallel to the communications model developed by Shannon in which the power *P* of the communication signal is absorbed into and then dissipates from the receiver while it selects a particular message [14]. The receiver must also handle additional energy caused by thermal noise *N* added to the power. The *P/N* ratio is called the ‘signal-to-noise’ ratio, but this term is not appropriate for EcoRI since there is no external signal. However, in both models there are the two energies dissipated in time, *P* and *N*, and there is a selection of specific states. Of course, a sufficiently strong thermal noise will eventually dislodge EcoRI from its specific binding, but this reversal is not the selection process we are interested in. Having set up these concepts allows us to apply powerful theorems from information theory to the recognition problem [7, 12, 15, 16].

However, the problem of recognition cannot be explained by thinking about the protein-DNA contact as a single interaction. Instead, there are multiple interactions including hydrogen, van der Waals and electrostatic bonds. To describe this set of interactions takes a series of numbers. Some of these interactions could be independent like the pins in a lock. As in a lock, it is advantageous for the pins to be independent because that way the lock can represent more combinations and is more secure [17]. Unlike a lock, the microscopic EcoRI molecule is continuously impacted and violently jostled by thermal noise (*N*). Each molecular ‘pin’ has a velocity that is the sum of many small impacts, so the central limit theorem from statistics tells us that the velocity will be approximately Gaussian [18]. So the moving parts of a molecule–the ‘pins’– that help EcoRI select GAATTC can be modeled as a set of independent molecular oscillators moving under the influence of thermal noise [7].

EcoRI potentially has many pins, each with a particular velocity. When one has a set of independent numbers they can be described as a point in a high dimensional space. Furthermore, when two independent Gaussian distributions are combined at right angles to represent their independence, the resulting 2-dimensional distribution is circular [19]. With three independent Gaussian distributions the combined distribution is a sphere and when there are more than three the distribution is still spherical, a hypersphere. The radius of the sphere is proportional to the square root of the thermal noise impacting on the molecule [7].

The higher the dimension of the sphere, the more the distribution converges to a single radius and the sphere skin or ‘thickness’ becomes smaller [7]. To see this, consider the volume of a ball (the region enclosed by a sphere) embedded in a *D* dimensional space,

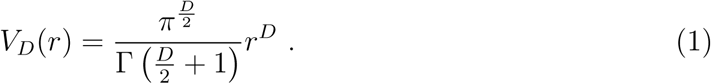

For a radius *r*, let half the volume lie in the shell between *r*_*_ *< r* and *r*, that is 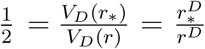. Rearranging gives 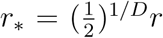 and as *D* → ∞, *r*_*_ → *r*, so the volume is densest near the surface.

So the state of EcoRI bound to GAATTC after dissipation can be represented as a hypersphere. Because the pins of EcoRI have an instantaneous position and velocity, at any one instant EcoRI is at a particular point on the sphere and moves by Brownian motion across the sphere surface. EcoRI bound to a different DNA sequence, such as CAATTC, is on a different hypersphere. If these two spheres were to intersect, then EcoRI would be able to bind sites other than GAATTC and this error would be fatal to the bacterium whose DNA is only protected at GAATTC. Thus the hyperspheres should not intersect. When EcoRI binds to DNA, that provides a finite amount energy that can be dissipated per binding (*P*), so there is a finite set of hyperspheres that can be bound. During evolution EcoRI will tend to minimize the binding energy, while the number of hypersphere states it selects between remains constant [12] so the hyperspheres become tightly packed together without intersecting (Fig 1). Thus EcoRI can evolve to bind efficiently, using the minimum energy to select between the maximum number of binding states. In addition, by using many interactions in a high dimensional space, the hyperspheres become sharply defined because the distribution around the sphere radius (the thickness) becomes smaller [7]. This allows EcoRI to evolve to reduce the number of times it cuts the wrong sequence, giving it a low error rate.

**Fig 1.**
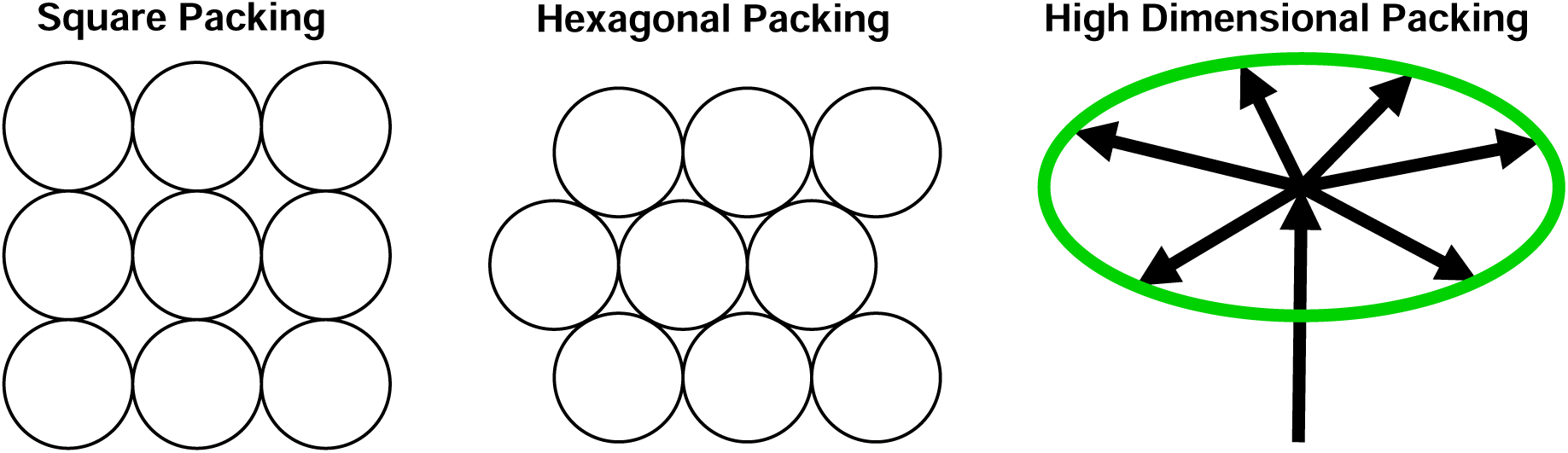
Sphere packing. Circles demonstrate square and hexagonal sphere packing in two dimensions. The hexagonal packing is 12% more dense. In higher dimensional spaces sphere packing is less intuitive. When hyperspheres pack together there is an odd property diagramed on the right side of the figure (which is derived from Shannon’s proof of the channel capacity theorem, Theorem 2 in his figure 5 [15]). The vertical arrow represents moving from the center of one hypersphere to the center of a second hypersphere. For Shannon, working with electrical communications, this voltage is proportional to the square root of the power dissipation, 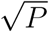. In a 100 dimensional space, the thermal noise in the second sphere (green circle) disturbs the signal in all directions, shown by splayed arrows with lengths proportional to 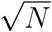. However, 99 of those dimensions do not perturb in the direction of the power dissipation. In his proof, Shannon neglected the 1% of the noise in the direction of the power since this represents the error, and it can be made as small as one may desire by increasing the dimensionality—in 1000 dimensions the error is only 0.1%. So relative to the direction of the power, the received hypersphere can be treated as a flat surface since all the other directions (splayed arrows) are at right angles to the power direction. If two hyperspheres are to be separated with as low an error as desired, then the power to get from one to the next must just exceed the thermal noise power of the first sphere, so 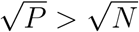 and *P > N*.

Shannon dealt with a closely related problem regarding maximizing the information that could be sent over a phone line for a given power (*P*, joules per second) [14, 15]. Messages from a transmitter can be broken into a series of independent voltages and so the set of numbers describing a particular message can be represented as a point in a high dimensional space which we call the ‘coding space’ since the message is represented by a code, the set of voltage values along each dimension. In addition, thermal noise on the phone wire causes the received voltage pulses to vary according to a Gaussian distribution. So if a message were repeated many times the received message points would form a sphere in the high dimensional space. When the receiver gets one of these noise-disturbed points, it can determine which of the possible transmitted messages is closest and thereby ‘decode’ the message to produce a clear noise-free signal for the person. Shannon recognized that the received spheres should not intersect if the receiver is to avoid ambiguity in decoding.

The receiver in a communications system selects particular symbols from all possible symbols that the transmitter might send. Similarly, a molecule such as EcoRI selects a particular state (binding to GAATTC) from an array of possible states (binding to any arbitrary 6 base long sequence). This concept applies to many other biological macromolecules. These ‘molecular machines’ include proteins that bind DNA, proteins that detect light such as rhodopsin in the eye and proteins that cause motion such as myosin moving on actin in muscle [7]. In every case the molecular machines dissipate energy in order to settle into one of several possible lower energy states and they do this despite the presence of violent thermal noise.

This paper answers the question of why restriction enzymes have such high fidelity despite being disturbed by thermal noise by showing how to calculate the coding space dimensionality of nucleic-acid recognizing molecular machines. The measured dimensionalities imply that restriction enzymes have evolved to exploit coding techniques only recently developed for modern communication systems. This in turn suggests that humans should also be able to build nanometer scale molecules that decode signals.

Background concepts important for understanding these results are basic information theory [20, 21], general molecular biology [22], and how to measure the information content of binding sites on DNA or RNA in bits [23,24] as in sequence logos [25]. In addition, messages in a communications system and the states of molecular machines can be represented by spheres packing together in a high dimensional coding space [7, 15, 26]. The isothermal efficiency is described in [12, 27]. For reviews, see [27–29].

The organization of this paper is as follows. Section 2 develops an equation for a lower bound on the dimensionality and we show how it applies to restriction enzymes in Section 3. Section 4 then develops an upper bound, and Section 5 shows that these bounds converge to a unique dimension, twice the number of bits. In Section 6 we analyze the dimensionality of over 4000 restriction enzymes and find that the 4- and 6-base cutting restriction enzymes prefer to operate in 16 and 24 dimensions. These dimensions contain dense hypersphere packings. Section 7 discusses the implications of restriction enzymes using good sphere packings and examines possible biological reasons why some restriction enzymes use packings in other dimensions. Section 8 examines biophysical mechanisms that restriction enzymes might use to attain high dimensional coding. Section 9 explores the relationship between high dimensional codes and restriction enzymes and examines additional details of how restriction enzymes recognize DNA in different dimensions. In Section 10 we discuss how the coding space may represent the biological fitness landscape through which restriction enzymes evolve. Finally in Section 11 we apply the theory to transcription factors in general and find that they are probably functioning not only in a high dimensional space, but the space dimensionality is not an integer so (by definition) they function in a high dimensional fractal space.

## 2 A lower bound on the dimensionality of molecular machines

Molecular machines are molecules that select specific states while dissipating energy [7]. The information, in bits, that a molecular machine can gain is the base 2 logarithm of the number of states it selects amongst [23–25]. The maximum number of bits that can be gained for the energy dissipated in a communications system is the channel capacity [15]. For molecular machines, we call the corresponding measure the molecular machine capacity [7]. Formulas for the capacities contain a term that represents the dimensionality of the coding space in which the state spheres are packed. Therefore a rearrangement of the formula leads to an equation for the dimensionality. This provides a step towards understanding the nature of the coding space of molecular machines.

The maximum number of distinct choices that a molecular machine can make in the presence of thermal noise *N* by dissipating energy *P* depends on these two factors and also on the number of independent moving parts of the machine or ‘pins,’ *d*_space_, following the lock and key analogy of molecular machines [7]. In communications, the channel capacity sets the upper bound on the rate that information can be faithfully transmitted [14, 15]. Corresponding to the channel capacity of communications systems a molecular machine’s capacity is:

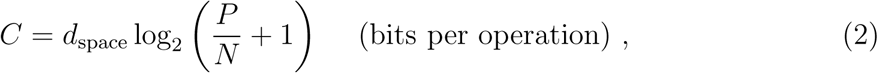

where a molecular machine operation is, for example, the process of going from non-specific to specific DNA binding by a nucleic acid recognizer [7]. This formula was derived by counting the maximum number of distinct molecular states, represented as spheres in a high dimensional space (see [27–29] for reviews), assuming white Gaussian noise. At the molecular level, relevant to the functionality of molecular machines such as restriction enzymes, the noise is overwhelmingly thermal, justifying the use of Shannon’s results. In order to faithfully transmit information, Shannon’s channel capacity theorem [15] implies that the sequence information a nucleic acid recognizing molecular machine uses to locate its binding sites, *R* = *R*_sequence_ [23], can evolve up to but not beyond this capacity:

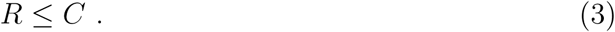

For nucleic acid recognizers, *R*_sequence_ is the area under a sequence logo [25]. The dimensionality of the coding space used to describe these states is:

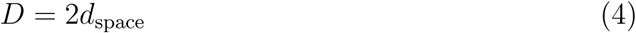

since there are both a phase and an amplitude for each of the independent oscillator pins that describe the motions of a molecule at thermal equilibrium [7]. Combining equations (2), (3), and (4) gives a lower bound for the dimensionality:

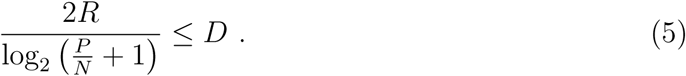

This lower bound is a function of the information gain *R* and the *P/N* ratio.

## 3 Applying the dimensional lower bound to restriction enzyme coding space

The maximum theoretical isothermal efficiency of a molecular machine is defined entirely by the dissipated energy *P* and the thermal noise *N* in terms of the normalized energy dissipation *ρ* = *P/N*:

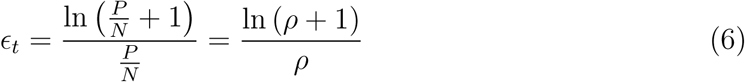

where 0 *< ϵ*_*t*_ ≤ 1 [12]. This expression was first used to describe the efficiency of satellite communications in terms of the ‘signal-to-noise’ ratio, *P/N* [30].

The efficiency of EcoRI and other molecular machines is observed to be close to 70% [12]. This can be explained if *P* ≈ *N*, in which case equation (6) shows *ϵ*_*t*_ ≈ ln 2 *≈* 0.69. The relationship between *P* and *N* measures the distance between hyper-spheres (Fig 1) so *P* = *N* implies that the state of being bound to one sequence is distinct from the state of being bound to a different sequence.

When there is a choice to be made among several molecular states, such as the strong discrimination EcoRI makes between GAATTC and single base changes of that sequence [5], then the molecular machine operates under the condition that its states are separated, which has been shown geometrically to be equivalent to

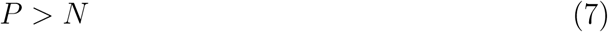

(Fig 1) [12]. This inequality limits the efficiency to 70%. Substituting into equation (5), the inequality (7) implies that

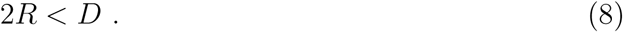

For fully evolved bistate molecular machines the dimensionality of the coding space is more than twice the information content of a binding site when the latter is expressed in bits. Thus, a lower bound of the dimensionality for the EcoRI coding space is found by noting that GAATTC is 6 bases or 12 bits, so EcoRI operates in a coding space of at least 24 dimensions. Similarly, as a consequence of the inequality (7), *d*_space_ supplies a lower bound for the channel capacity in equation (2),

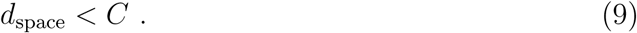

For example, given *P > N*, if *P* = 31*N* then *C* = 5*d*_space_, so *C > d*_space_.

## 4 An upper bound on the dimensionality of molecular machines

The higher the dimension that a molecular machine can work in, the more the probability density tightens around the radius of the hyperspheres [7]. This suggests that biological systems may tend to evolve to extremely high dimensions to reduce the error rate caused by switching between the hyperspheres. So having determined a lower bound on the dimensionality of a molecular machine (equation (5)) is tantalizing but unsatisfying because biological systems may have much higher dimensionality. For this reason we sought an upper bound on the dimensionality.

The dimensionality of a molecule is related to the number of degrees of freedom (*ν*) that a molecule has. For *n* atoms there are 3 independent axes each atom can move on, but the three translational motions and three rotations about the axes do not contribute to the functioning of the machine, so there are only

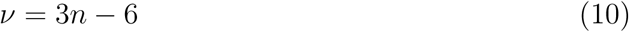

degrees of freedom. For water *n* = 3 so *ν* = 3 normal modes that can be observed in the vibrational spectrum of the molecule. These three motions can be described by common arm exercises with the head representing oxygen and the fists hydrogen: pushup/pullup (*ν*_1_ symmetric stretch), jumping jack (*ν*_2_ bending mode) and one-two punch (*ν*_3_ asymmetric stretch) [31].

Although the number of degrees of freedom of an entire molecule consisting of *n* atoms is 3*n* − 6, the relevant number of degrees of freedom involved in the molecular machine selection process coding space (*D*) is most likely much smaller [7]:

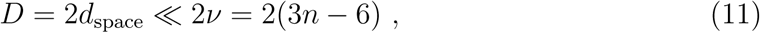

because to be able to evolve each molecular machine ‘pin’ consists of an average of up to *n/d*_space_ ≫ 1 atoms. For a large molecule like EcoRI with thousands of atoms, the relevant degrees of freedom (*D*) for DNA binding will be much smaller than given by equation (11), so that relationship does not give a useful upper bound.

As Jaynes pointed out [32,33], based on the classical equipartition theorem, the energy per degree of freedom of a single thermal oscillator in a molecular machine (lock pin) is 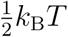; with *D* degrees of freedom the total thermal noise flowing through a molecule during one dissipation step of *P* that selects a specific molecular state is

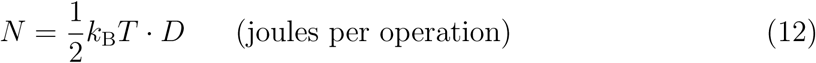

(see also equation (31) in [7]).

For molecules that make distinct decisions by selecting between nonoverlapping hyper-spherical states, the inequality (7) applies. Substituting (7) into equation (12),

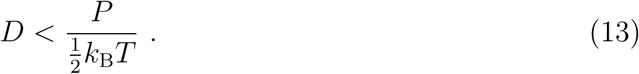

This provides a upper bound on the functional dimensionality, whereas equation (5) provides a lower bound.

We convert equation (13) to a more useful form by noting that the energy available in coding space for making selections at one temperature and pressure [7, 12, 27] is the Gibbs free energy:

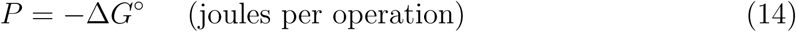

[12]. The maximum number of bits that can be gained for that free energy dissipation is

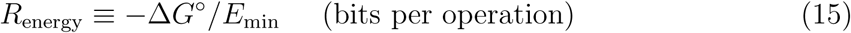

[12]. *E*_min_ can be derived from information theory [34] or the second law of thermodynamics [16, 35]. It serves as an ideal conversion factor between energy and bits:

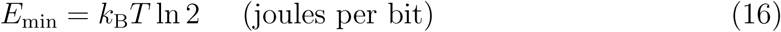

[12]. Further, a ‘real’ isothermal efficiency *ϵ*_*r*_, that may be less than the theoretical efficiency of equation (6),

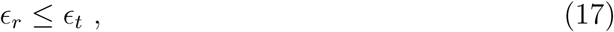

can be measured by the information gained, *R, versus* the information that could have been gained for the given energy dissipation, *R*_energy_:

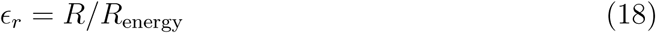

[12]. Successively combining equations (14) to (18) gives

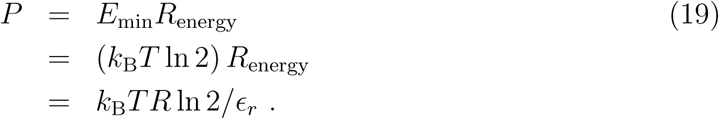

Inserting this result into equation (13) gives

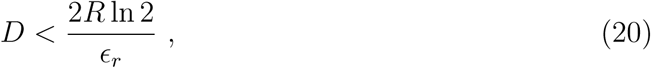

which we recognize as an upper bound on the coding space dimensionality as a function of the information gain *R* and the isothermal efficiency *ϵ*_*r*_.

## 5 Pincers on the dimensionality of molecular machines

Having determined both a lower bound (equation (5)) and an upper bound (equation 20) on the dimensionality of the coding space, we have the opportunity to determine what will happen as the molecular machine evolves to be optimally efficient.

Combining equations (5) and (20) gives

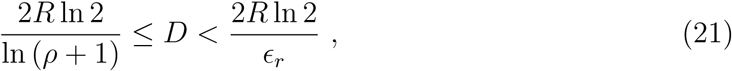

where we have recast the left hand side in terms of *ρ* = *P/N* and expressed the logarithm in base *e* to emphasize the striking symmetry of the two sides (the numerators are identical and the denominators are related by equations (6) and (17)). The dimensionality is constrained to lie between these bounds, which form closing ‘pincers’ as the molecular machine evolves to become optimal. In the limit as *P* → *N* and *ϵ*_*r*_ evolves to its maximum value of ln 2 [12], both sides converge to 2*R*, and *D* is squeezed between them. An optimally evolved molecular machine will operate in 2*R* dimensions:

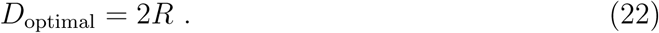

However, because the state hyperspheres have a finite thickness [7], *P* must at least slightly exceed *N*, and the left hand side of (21) remains slightly smaller than 2*R*. Likewise, because of equations (6) and (17), *P > N* also means that the efficiency *ϵ*_*r*_ is slightly below ln 2, which makes the right hand side of (21) sightly larger than 2*R*. So the dimensionality is restricted to a small interval. This allowed variation in the coding space dimensionality is caused by the effective thickness of the sphere surface fuzziness (which depends on the dimensionality *D* itself) and the evolutionarily acceptable error rate determined by the environment which limits how closely the coding spheres can approach each other and still allow survival [7].

For example, DNA polymerase has a certain error rate, and of course if that error rate were to increase the organism would experience a higher number mutations and be at a selective disadvantage. However, there are also mutations of the polymerase that decrease its error rate [36]. This would reduce the mutation load on the organism, but presumably it does not occur in the wild because the organism would then be less able to evolutionarily adapt to changing conditions compared to siblings that have the higher error rate. In many but not all cases they would also replicate more slowly. Likewise, the error rate for translation is about 1 in 1000 amino acids [37] which means that roughly one in every three proteins has an error. Yet organisms survive quite well at this error rate. The error rate set by the environment of the organism in turn determines the acceptable placement and thickness of the spheres.

Note that the lower bound constraint comes from equation (3), the channel capacity theorem of Shannon [15], which limits the efficiency *ϵ*_*r*_. The upper bound comes from equation (7), the finite energy available to perform state selections (*P*) relative to the thermal noise (*N*), which satisfies the biologically required separation of molecular states [12]. These two independent bounds are plotted on a graph of the efficiency curve (Fig 2, equation (6) [12]).

**Fig 2.**
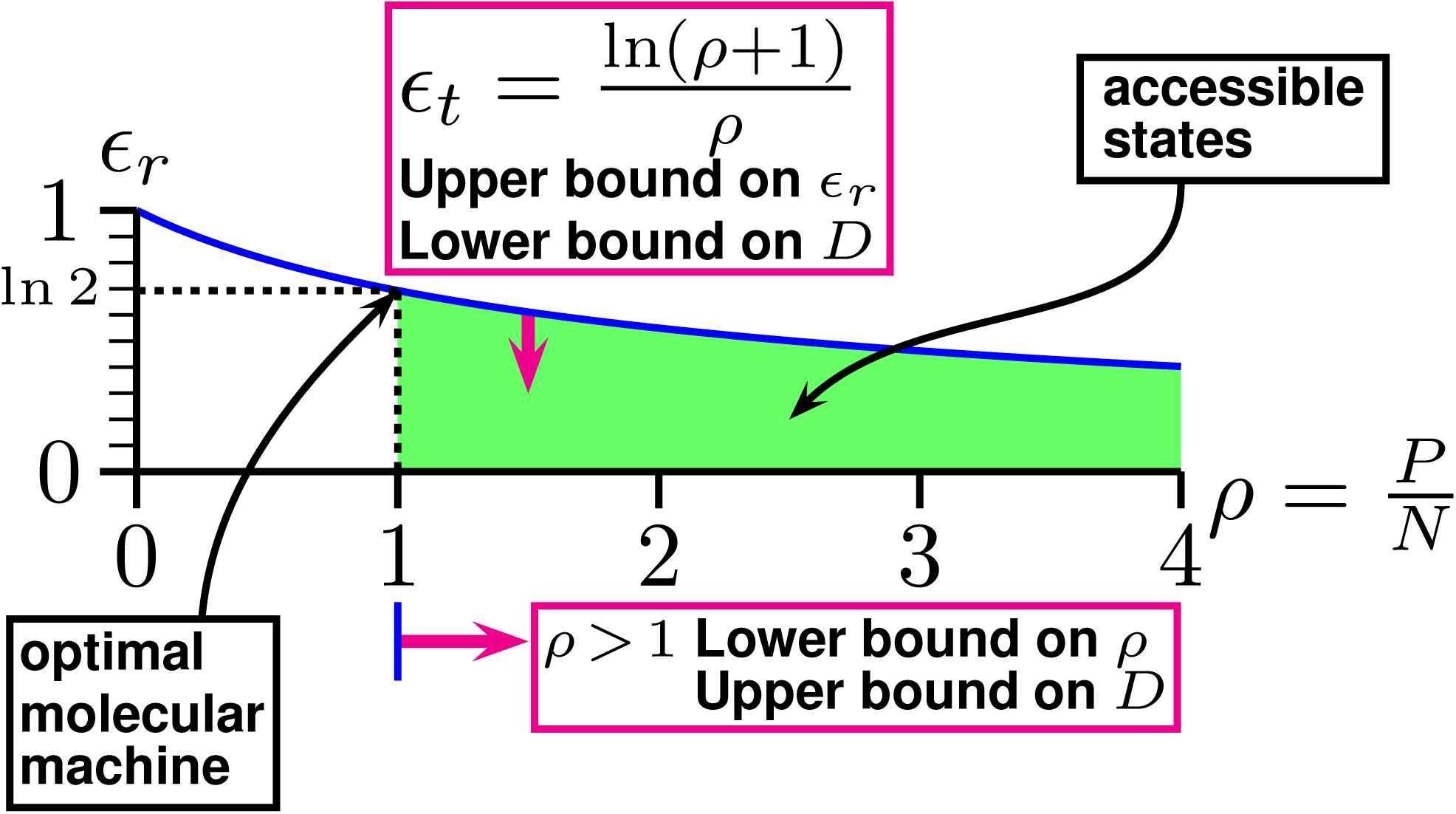
Isothermal efficiency curve for molecular machines showing bounds that constrain the coding space dimensionality *D*. Real molecular machines that select between two or more distinct states may have parameters anywhere in the shaded (green) area in which the real isothermal efficiency *ϵ*_*r*_ is bounded above by the theoretical isothermal efficiency *ϵ*_*t*_ (equation (6)) and to the left by the power to noise ratio *ρ* = *P/N >* 1 (equation (7)). During evolution, they tend to lose unnecessary energy dissipation, which decreases *ρ* towards the lower limit of *ρ* = 1. Independently, they tend to increase their information use (*R*) for the energy dissipated, which increases *ϵ*_*r*_ toward the theoretical maximum *ϵ*_*t*_ determined by the channel capacity. These factors lead to an ‘optimal’ molecular machine in which *ρ* = 1 and *ϵ*_*r*_ = *ϵ*_*t*_ = ln 2. At that point the dimensionality has been squeezed in a pincers (equation (21)) until it reaches *D* = 2*R*.

The efficiency curve is an upper bound representing functioning at the channel capacity (equation (3)). Points below the curve have *R < C*. Since the dimensionality parameter *D* = 2*d*_space_ is part of the upper bound for *C* in equation (2), this leads to the lower bound on *D* in equation (5). Independently of that, the *ρ* = *P/N >* 1 ratio on the horizontal axis of Fig 2 is orthogonal to the efficiency and channel capacity of the vertical axis. The thermal noise *N* is determined by the absolute temperature of the molecular machine and the dimensionality (equation (12)). Since *P* is an upper bound on the noise, equation (7) leads to an upper bound on the dimensionality in equation (20).

These two independent constraints on *ϵ*_*r*_ and *ρ* determine the possible range of the dimensionality. As shown previously [12], the normalized energy dissipation *ρ* will tend to decrease over evolutionary time because excess contacts that are not required for maintaining information will be lost by mutation. This decrease will continue until at *ρ* = 1 the distinctness of molecular states is threatened by a large error rate caused by the increasing intersection of after state spheres. Meanwhile, the efficiency *ϵ*_*r*_ will tend to increase up to ln 2, squeezing the dimensionality towards a single value, 2*R* by equation (21). This result is consistent with the dimensional analysis of Collier [38], who showed that information is related to the degrees of freedom, which is of course the dimensionality of the space.

## 6 Dimensionality of restriction enzyme coding space

Now that we know that the dimensionality of an optimal molecular machine is simply twice the number of bits that it selects amongst (equation (22)) we can determine the dimensionalities of the thousands of known restriction enzymes if we assume that they too are optimal. In the case of restriction enzyme EcoRI, experimental data [13] show that the efficiency is close to *ϵ*_*r*_ = ln 2 [12], so that *ρ* = *P/N* must be close to 1. Thus, although complete data are not available about specific and non-specific binding of other restriction enzymes, we assume that, like EcoRI, these molecular machines have also evolved to be close to ln 2 efficiency as shown in Fig 2. We believe this is a reasonable assumption because they should all distinguish their binding sites from other sequences with as small an error as possible so their hypersphere states should not overlap significantly (*P > N*), and they should maximize the information gain for energy dissipation (the efficiency) but they cannot exceed the efficiency upper bound shown in equation (6) [12, 27].

Since a 6 base cutting restriction enzyme recognizes *R* = 6 × 2 bits = 12 bits, from equation (21), we see that the EcoRI coding space must be close to 24 dimensions. If we characterize a restriction enzyme by the number of bases it recognizes, then there are a maximum of two bits per base, so the dimensionality of a 70% efficient molecule is:

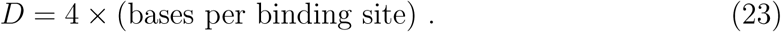

Thus, EcoRI, which has a 6 base recognition site GAATTC, works in 24 dimensions, while TaqI, which recognizes only the 4 bases TCGA, should work in 16 dimensions. The highest known dimension used by a restriction enzyme is 32 dimensions for restriction enzymes such as NotI (GCGGCCGC) and SfiI (GGCCNNNNNGGCC), which cut DNA at patterns 8 base pairs long [39, 40].

In the case of restriction enzymes that digest at partially variable patterns such as GT(T/C)(A/G)AC (HincII), we can use the information needed to describe the pattern to predict the dimension. In this case, for the first two and last two bases (GT and AC), a total of 8 bits are required, while for each of the middle two bases only 1 bit is required to distinguish two of the four bases so *R* = 10 bits per site [23], even though the binding site is 6 bases long. Thus, if HincII is optimal at 70% efficiency, it should operate in 2*R* = 20 dimensions. In the special case where a base is avoided by a restriction enzyme, we record the information as 2 − log2 3 bits for that base [23]. So formula (23) has only a limited application. The number of bits in a binding site is not strictly computed from the physical length of the site, but rather from the average number of bases if all the information were compressed into the smallest region possible [23]. This is the ‘area’ under a sequence logo [25].

The predicted dimensionality of over 4000 Type II restriction enzymes in Roberts’ REBASE database [40] is given in Table 1 and Fig 3A. There are two major peaks at 24 and 16 dimensions, corresponding to 6 and 4 base cutters. There is also a minor peak at 32 dimensions for the 8 base cutters.

**Table 1.**
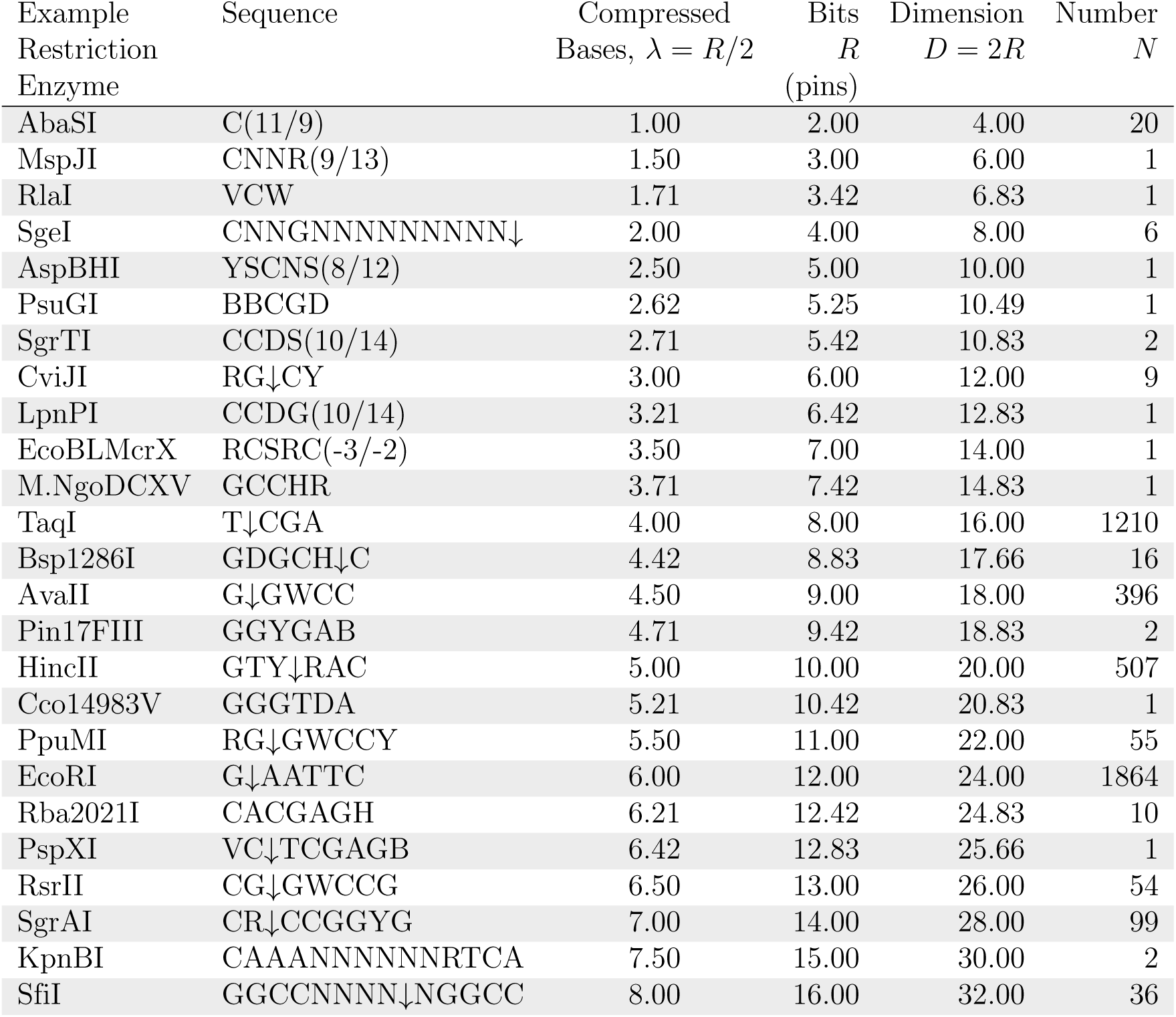
Coding space dimensionality (*D*) and number (*N*) of restriction enzymes. The information content in bits, *R*, of the recognition sequence of 4297 restriction enzymes from REBASE (restriction enzyme database) http://rebase.neb.com or ftp://ftp.neb.com/pub/rebase/ version allenz.801 (Dec 27 2017) [40] was computed. A fully conserved base (A, C, G, T) contributes 2 − log2 1 = 2 bits, two possibilities (R=G/A, Y=C/T, M=A/C, K=G/T, S=C/G, W=A/T) contributes 2 − log2 2 = 1 bit, three possibilities (B=C/G/T, D=A/G/T, H=A/C/T, V=A/C/G) contributes 2 − log2 3 ≈ 0.42 bits and any allowed base (N) contributes 2 − log2 4 = 0 bits [23, 41]. The sum of the information at each base, *R*, was used to find the corresponding number of compressed bases (*λ* = *R/*2) and then the coding dimension (*D* = 2*R*), assuming that each enzyme has an efficiency of *ϵ*_*r*_ = ln 2 and *ρ* = 1 so that there is a unique dimension according to equation (21). The most commercially available enzymes and their reported recognition sequences are given as examples. When the DNA backbone cleavage site is known it is indicated by an arrow (↓). The distance to cleavage sites outside the given sequence is shown in parenthesis for the corresponding and complementary strands. Star activity (variation within the canonical site) and flanking sequence effects are found for many restriction enzymes [42]. However, the patterns in the database are reported as consensus sequences that may distort the information content [43], and so may affect the results given here.

**Fig 3.**
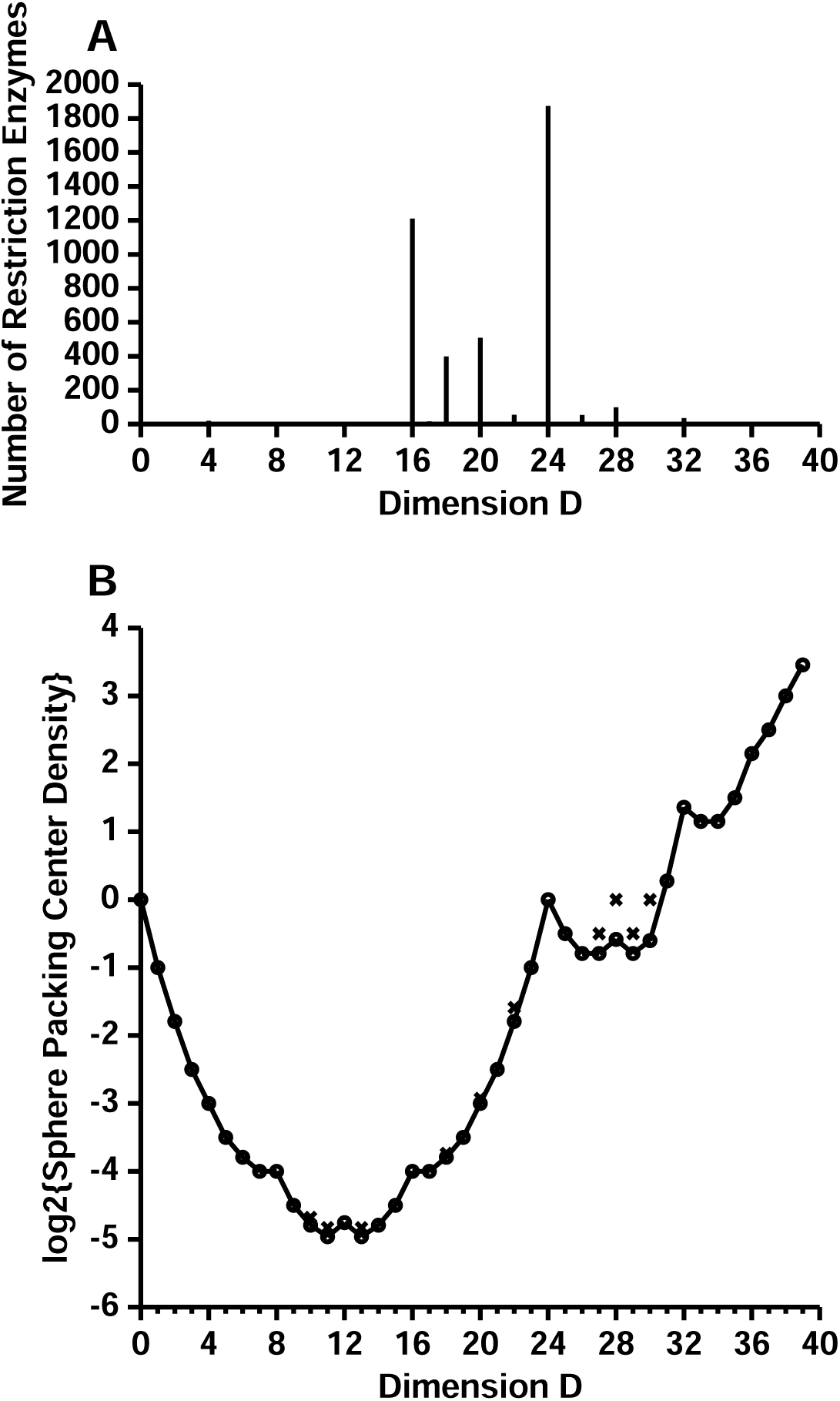
Comparison of restriction enzyme frequency and best known sphere packing density in different dimensions. A. Coding dimensions used by restriction enzymes. The number of enzymes at each dimensionality is plotted from Table 1. B. Best known sphere packings in high dimensions were given by Conway and Sloane [26,44]. The graph is equivalent to their Figure 1.5; see Table I.1(a), Table I.1(b) on pages xix and xx; and pages 14 to 16. The updated sphere center density formulas used here were from http://www.math.rwth-aachen.de/~Gabriele.Nebe/LATTICES/density.html (Last modified Feb. 2012, accessed Jan 06, 2018). The sphere center density, *δ*, is the number of sphere centers per unit volume when sphere radii are set to 1. Without the logarithm, a graph of *δ* versus *D* appears nearly flat from *D* = 7 to *D* = 18. Circles (○) represent lattice packings; x’s (×) represent nonlattice packings.

The 4297 enzymes we studied consist mostly (98%) of Type II restriction enzymes. A set of 482 Type I enzymes, which cut randomly at a distance away from where they bind specifically, were also collected from REBASE [40] (http://rebase.neb.com/rebase/rebadvsearch.html on 9/30/2019 using the search paramters: ‘Type I’, ‘Specificity subunit’, ‘Non-putative’, ‘Prototype’ and columns: ‘Enzyme name’ and ‘Recognition sequence’). They primarily use 24 (105 cases), 26 (112 cases) and 28 (206 cases) dimensions. Since the additional enzymes are a small fraction of the database our conclusions would not change by including them.

## 7 Biological lattices in high dimensional spaces explain restriction enzyme 4 and 6 base pair preferences

Ever since Shannon published his theory of hypersphere packing as a description of a communications system [15] mathematicians and engineers have been determining how best to pack spheres together in high dimensional spaces [26]. The restriction enzymes appear to favor particular dimensions for their coding spaces, so we can compare their preferences to the best known packings that humans have determined.

In two dimensions there are two ways to regularly pack circles: in a square lattice or in a hexagonal lattice (Fig 1). The square packing fills Δ = *πr*^2^*/*(2 × *r*)^2^ = 79% of the plane, while a hexagonal packing fills 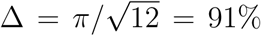 [26]. Hexagonal packing is more dense than square packing. In general, the sphere packing density is

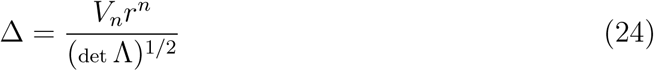

where *V*_n_ is the volume of an *n* dimensional sphere with radius *r* [45–47], and det Λ is the determinant of the lattice Λ. The determinant provides the volume of the polytope that the sphere is encased in. For convenience, Leech introduced the concept of the sphere center density

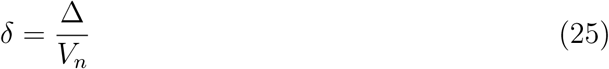

which counts the average number of sphere centers per unit volume of the space [6]. Leech, Conway and Sloane rescaled or normalized the sphere center density in several different ways to emphasize the symmetries of the sphere center density as a function of dimension [6, 26, 48]. As these rescalings do not have any biological significance that we are aware of, we only take the logarithm of *δ* to graph the best known sphere packings up to 40 dimensions (Fig 3B). Since each sphere center represents one sphere, and in a biological context spheres represent biological states, the center density is a measure of the number of states available to the system.

Shannon recognized that the packing of spheres in higher dimensions corresponds to the problem of faithfully transmitting a series of distinct messages over a noisy communications channel [15, 26, 49, 50]. In this model, transmitted messages are points in a high dimensional space, and each sphere represents a message received with Gaussian noise added along each dimension. Most of the sphere density is on the surface in high dimensions [7]. To avoid message ambiguity, the spheres must not intersect, which spaces the transmitted messages and allows a decoding that removes the noise from the received signal. The total volume available in which to pack spheres is a large sphere whose radius is determined by the power and thermal noise absorbed by the receiver, while the volume of a smaller message sphere is determined by the thermal noise alone [15].

A corresponding theory also describes the states of molecular machines as spheres in high dimensional space [7]. In both theories, the maximum number of possible messages (or molecular states), known as the capacity, is determined from the number of small spheres that can be packed together inside the larger sphere. As in 2 dimensions, there are many possible ways to pack high dimensional spheres; the more spheres that can be packed together, the more the channel can be utilized. Because of its application to communications and a variety of mathematical and physics fields, the highest sphere packing densities in various dimensions have been determined, as shown in Fig 3B.

The histogram in Fig 3A is compiled from data corresponding to 4297 restriction enzymes, whereas the center density graph in Fig 3B is an exact mathematical result for 51 lattice packings. It is striking how well they match at *D* = 16 and *D* = 24, corresponding to 4 and 6 base cutters. The histogram also has minor peaks at *D* = 18 and *D* = 20 that are not reflected in the density plot. We argue below that this happens for biological reasons. Similarly, the peak at *D* = 32 in the center density plot does not have a corresponding large peak in the histogram. As discussed below, we explain that this is for biological reasons as evolution seems to prefer recognition and excision of smaller pieces of DNA.

The most dense known sphere packing is in 24 dimensions, a packing known as the Leech lattice (symbolized as Λ_24_) [6,26,51–57]. This packing has been extensively studied, and because of its density it was used in a commercial Motorola modem [58]. Surprisingly, the most common dimensionality of the restriction enzymes is also 24 dimensions, as shown in Fig 3A. This suggests that EcoRI and the other restriction enzymes may be 6 base cutters because that takes advantage of the dense packing of the Leech lattice. In other words, we hypothesize that the reason so many restriction enzymes are 6 base cutters is that they have discovered the Leech lattice packing by Darwinian evolution. How the Leech lattice is implemented by the atomic structure of restriction enzyme proteins is not known.

For restriction enzymes, the next most commonly used dimensionality is 16 dimensions (Fig 3A), and we see in Fig 3B that a good packing, the Barnes–Wall lattice Λ_16_ (BW_16_), has also been found for this dimension relative to the other dimensions of similar magnitude [44]. There are an estimated 10^7^ good packings equivalent to BW_16_ [59]. Thus, the 4 base cutting restriction enzymes may be using 8 bit recognition to take advantage of the good hypersphere packings possible in 16 dimensions.

The longest known restriction enzyme sites have 8 bases, and so these enzymes should use a 8 × 4 = 32 dimensional space. Correspondingly, 32 dimensions also represents a peak in the known dense lattices called the Quebbemann’s lattice, *Q*_32_ [44]. Cohn and Elkies report a peak in 28 dimensions [52] and, intriguingly, this corresponds to 99 cases of 7 base cutters (Table 1). Transcription factors in *E. coli* have information contents in the range 16 to 23 bits [23]; these may function with the high density packings known to exist above 40 dimensions.

On the lower dimensional end, it is worth noting that there is a small local maximum for the density of sphere packings at 12 dimensions. This would correspond to a 3 base long biological object for which the obvious candidate is the codon of the genetic code. It may be that the genetic code functions in a 12 dimensional space, but the coding is probably not performed via the cubic lattice *Z*^12^ suggested by Sadegh–Zadeh [60] since there are better packings such as the Coxeter–Todd lattice *K*_12_ [26]. Finally, in 8 dimensions the most efficient possible sphere packing is on an *E*_8_ lattice [56]; this corresponds to 2 base pair recognition. Biologically this code might be used for precisely recognizing and methylating CpG base pairs, the basis of an important epigenetic control [61, 62]. Thus, all of the peaks in Fig 3b could correspond to known biological systems.

Restriction enzymes can evolve from one dimension to the next since this only requires increasing or decreasing the number of base contacts [63, 64]. So there is some fluidity in the dimensions chosen, but for several reasons we do not expect a complete correspondence between Fig 3A and Fig 3B. First, because restriction enzymes have evolved, there is a good deal of history in the current choices and some of this may be locked in. Some patterns will be common simply because that particular bacterial species is prevalent and their restriction enzymes were discovered more easily than others. Second, the 8 base enzymes (32 dimensions) won’t attack an invading DNA as frequently as shorter ones, so bacteria may tend to avoid using higher dimensions. Third, short patterns that cut frequently would necessitate more self-protective methylation and so would be expensive [64]. Fourth, evolutionary pressures to increase or decrease the binding site information may not be equal [64]. Finally, unknown effects could come into play to eliminate, for example, most of the 5.5 base (22 dimensional) restriction enzymes even though the 5 (20 dimensional) and 6 (24 dimensional) base sites are quite common.

The information content of transcription factor DNA binding sites evolves based pre-dominantly on the size of the genome and the number of binding sites [23, 24]. Unlike transcription factors, the information content of restriction enzyme sites cannot evolve based on invading genomes because there are no regular specific sequences to bind to. However, the size of the intruding genome does provide some criterion since restriction enzymes protect bacteria from invading bacteriophage. Typical sizes are on the order of 40, 000 base pairs, such as *λ* (48502 base pairs) [65] and T7 (39937 base pairs) [66]. The restriction enzyme must cut the invader at least once, and preferably more, to disable the phage genome. Thus, it requires approximately log2 48502 = 16 bits in the site to attack *λ* once. A 12 bit (6 base pairs) site such as EcoRI would cut *λ* 2^16−12^ = 16 times; 5 sites are observed. Perhaps this number is lower than expected because phage evolve away from restriction sites; EcoRI would cut T7 8 times but none are observed. An 8 bit (4 base pairs) site would cut more frequently than a 12 bit site but the cell would then have to methylate 2^12−8^ = 16 times as many sites. Perhaps this is one reason that 6 base restriction sites are more abundant than 4 base cutters: 6 bases is short enough that phage are killed but also sufficiently long that methylation is minimized.

Only the best choices of sphere packings in biologically useful dimensions may be reflected in the restriction enzymes. The central suggestion of this paper is that, although restriction enzymes are highly divergent [67–71], most of them have discovered that sphere packing in 16 and 24 dimensions is more dense than packings in other dimensions. This provides an explanation for why 4 and 6 base cutters have been found so frequently. In addition, the significant peaks at 18 and 20 dimensions in Fig 3A suggest the biological use of dense codes in those dimensions that may be consistent with known packings.

We regard an explanation as being a theoretical justification for an observed feature within a data set. Alternative reasons for restriction enzyme 4 and 6 base preferences might including ease of coding or efficacy of protein folding, but there needs to be a specific hypothesis justified by a mathematical model in order to provide a competing explanation. To our knowledge, there isn’t an alternative model that explains the features we have noted. In our view, since restriction enzymes are highly divergent [67–71], ease of coding and protein folding are unlikely to explain the convergence to 4 and 6 bases. Also, our model provides a compelling explanation for which dimensions are preferred.

## 8 Mechanism of high dimensional coding

For an evolved molecular machine *D* = 2*d*_space_ (from equation (4)) and *D* = 2*R* (from equation (21)) so *d*_space_ = *R*. So, curiously, for an optimal molecular machine the number of bits is the number of pins. However, how the independent pins are implemented in molecular architecture is a difficult open problem. As in genetics, the underlying mechanism of DNA recombination was not initially known but the results, linearity of genes, were still valid. Here, we know the dimensionality from the theory, but we would also like to know how the molecule works.

There are at least two basic mechanisms by which high dimensional coding could be implemented by molecules: direct contacts and vibrational modes. For example, EcoRI cuts double stranded DNA at the sequence GAATTC. In the co-crystal between EcoRI and this sequence, McClarin *et al.* [11,72] observed that each of the 6 bases is contacted with two hydrogen bonds, for a total of 12 specific hydrogen bonds. If each hydrogen bond corresponds to a single ‘pin’ of the molecular machine, with two degrees of freedom per pin [7], there would be 24 dimensions. That such contacts often act independently, and so could be coding space dimensions, is suggested by experiments on several other recognizers [7,73–77]. However, experiments with mutant EcoRI imply that it uses more than just hydrogen bonding in recognition [78], and bases of DNA recognition proteins are not entirely independent [79,80]. Though including dinucleotides may be sufficient [81,82], finding the important independent dimensions may be challenging. All such pairwise correlations can be displayed with a 3 dimensional sequence logo [83]. Alternatively, the coding space could consist of normal modes of molecular vibration since these are by definition independent [84]. In particular, localized vibrational modes called ‘discrete breathers’ [85] may represent the molecular machine pins.

## 9 Coding spaces

In classical information theory, a continuous communications signal, such as a song, can be represented by a series of independent numerical values [15]. An analog signal of duration *t* seconds that has a range of frequencies (bandwidth) *W* is described by *D* = 2*tW* Fourier components. Since these sine wave amplitudes are independent, they define *D* numbers and hence a single point in a *D* dimensional coding space. Because they are designed from scratch, the dimensionality in communications systems is known *a priori*. By contrast molecular systems, which also have been shown to use coding spaces [12], do not have a known dimensionality so determining this parameter is an important step towards fully characterizing and understanding their function.

Equations (5) and (20) establish lower and upper bounds on the dimensionality of molecular machines. These constraints can be represented geometrically (Fig 2). The restriction that the information *R* cannot be larger than the machine capacity *C* (equation (3)) ultimately comes from Shannon’s 1949 model of communication in which he divided the volume of a large ‘*before*’ sphere, representing the space of all possible messages, by the volume of a small ‘*after*’ sphere, representing a single message expanded in all possible directions by thermal noise, to determine the maximum number of possible distinct messages *M* in time *t* and hence the channel capacity in bits,

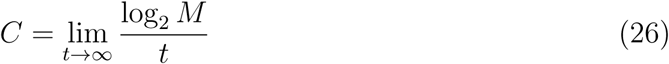

[15]. The corresponding model for molecular states (equation (2)) [7] leads to the lower bound on the dimensionality (equation (5)). In this case there are two geometrical constraints, state spheres must not intersect and the state spheres are confined to the larger sphere defined by the available energy. The observation of 70% efficient molecular machines comes from the restriction that for biological states to be distinct, the *after* state spheres must avoid intersecting each other [12], as expressed by *P > N* (equation (7)). The two constraints on the dimensionality therefore come from the *after* state spheres bumping into each other and from them being compressed within the larger *before* sphere.

We found that when a DNA binding protein evolves to be optimally efficient, the upper and lower dimensional bounds converge to twice the information content of the binding site as measured in bits (Fig 2). Using this result, we found that the common 6 base pair recognizing restriction enzymes, which require 12 bits to describe their pattern, use a 24 dimensional coding space. When EcoRI is bound to a DNA sequence its state can be described as a sphere in the high dimensional coding space with each of the possible 4^6^ = 4096 hexamer sequences represented by a different sphere [7]. If the sphere for EcoRI bound to GAATTC were to overlap with any other sequence sphere, then EcoRI could bind to and cut at inappropriate locations that are unprotected by the corresponding methylase, leading to death [4]. Since EcoRI binding to sequences other than GAATTC is at least 10^6^ fold down in digestion [5], these spheres effectively do not intersect. Excess binding energy that retains the same binding pattern will be lost by mutational changes in the EcoRI protein structure, so the spheres must be tightly packed together in the 24 dimensional space. Remarkably, it has already been shown by coding theorists that the best known sphere packing is the Leech lattice in 24 dimensions [26,44,57]. Likewise the 4 base pair restriction enzymes use a 16 dimensional coding space, and there are good packings known in that space (Fig 3) [59]. Thus, there is a correlation between commonly observed restriction enzyme DNA site sizes and the best packing of spheres in high dimensional spaces. Could this be a coincidence? We believe it is not for the following additional reasons.

First, the data sets are large. The entire collection represents nearly 4300 restriction enzyme sites (Table 1). Restriction enzymes are initially discovered by their ability to digest DNA, and this method does not indicate the sequence of the binding site, which is unknown until after the enzyme has been isolated, purified, and characterized. Odd classes of sites are noticed and publicized because these are eagerly sought as research reagents. Likewise, the data on different kinds of high dimensional sphere packings (Fig 3b) represent research efforts spanning the 70 years since Shannon’s publication in 1948, and there are strong economic incentives to discover and publicize new packings because they can be used to improve communications.

Second, the correlation between sphere packing and restriction enzymes was derived without introducing any free parameters to the boundary equations for the dimensionality. It is a natural consequence of previously established molecular machine theory [7, 12, 16, 27].

Presumably natural systems discovered the Leech lattice long ago, but the details of how a small protein can implement such a code are unknown. However, the fact that restriction enzymes have apparently discovered good codes should help us to understand how they can recognize short DNA sequences so precisely. Conversely, understanding the molecular mechanism of restriction enzyme decoding could lead to single-molecule communications devices [7].

The distribution of restriction enzyme choice of dimensionality is well explained by the best packings of spheres in various dimensions (Fig 3). The major peak of 6 base cutting restriction enzymes is most likely explained by their use of the Leech lattice in 6 × 4 = 24 dimensions. Likewise, the peaks at 16 and 32 dimensions correspond to the 4 and 8 base cutters respectively. In addition, restriction enzyme use of 18, 20, and 28 dimensions appears to correspond to good nonlattice packings that are known in those dimensions. This leaves three holes in the distribution at 22, 26, and 30 dimensions which are rarely used by restriction enzymes but which have decent sphere packings. We suggest that restriction enzymes with dimensions close to a peak evolve into the peak. For dimension 30, Table 1 gives the example of KpnBI with the recognition sequence CAAANNNNNNRTCA. Notably this is an asymmetric recognition sequence with a part that recognizes exactly 4 bases CAAA on the 5′ side and RTCA on the 3′ side. Recognition of a purine R is typically accomplished by a single hydrogen bond to the N7 position of either A or G [86,87]. If an additional contact or hydrogen bond into the major groove evolves (for example to the N6 of A or O6 of G and on the complementary strand the methyl of T, O4 of T or N4 of C), then the enzyme could specify exactly A or G and the dimensionality would increase to 32. This could improve the sphere packing according to our current knowledge of lattices in 30 and 32 dimensions, so the lack of *D* = 30 enzymes is likely to be because most have already evolved to the nearby better packing. Since there are two known *D* = 30 enzymes according to Table 1, we can test this idea by inspection of the other one. Indeed, that enzyme is Eco851I with recognition sequence GTCANNNNNNTGAY. Since Y pairs to R on the complementary strand, alteration of the terminal Y to a specific base would switch this enzyme from 30 to 32 dimensions by the same mechanism. Indeed, a similar explanation for an evolutionarily easy switch from 26 to the 24 dimensional Leech lattice is suggested by the enzyme RsrII CGGWCCG, while the 22 dimensional PpuMI RGGWCCY has three such opportunities. In each of these cases, merely having the disfavored dimension leads to a pattern vulnerable to evolution to a nearby dimension. Whether there are biological or mathematical constraints that prevent the rare cases from evolving to the best packing dimensions is unknown.

Two additional biological constraints on the distribution of restriction enzyme dimensionalities were mentioned earlier. We expect few if any restriction enzymes to function at low dimensionality (below 4 bases or 16 dimensions) since such enzymes would digest DNA frequently and so would require extensive methylation protection which may be disadvantageous. Restriction sites longer than 8 bases or 32 dimensions would only be found rarely on invading DNA and so presumably these too would not have much advantage. These factors limit the range of functionally useful dimensions.

## 10 Coding space as a fitness landscape

Considering how well the restriction enzyme frequency (Fig 3A) and best sphere packing center density (Fig 3B) distributions match overall given the biological constraints on the range that restriction enzymes can function in, the sphere packing center density distribution appears to be a measure of fitness for restriction enzymes evolving over a high dimensional adaptive fitness landscape, similar to the high dimensional spaces described by Wright [88–91].

For biological systems, the number of sphere centers corresponds to the number of distinct states the system can be in. The center density *δ* (equation (25)) is therefore a more appropriate measure than the filled volume of the lattice defined by Δ (equation (24)) since biological systems evolve to have distinct states [12]. The volume of the state is itself irrelevant. So we propose that log2 *δ* vs. *D* represents the biological coding landscape. Plotting log2 *δ* instead of *δ* emphasizes the detailed features of the curve. A biological system can evolve to obtain the highest number of distinct states by maximizing the sphere center packing density *δ*. Because the capacity is the logarithm of the number of states, this also maximizes the information and the efficiency.

Since the logarithm is monotonic, if *δ*_1_ *> δ*_2_, log2 *δ*_1_ *>* log2 *δ*_2_. Examining Fig 3B, we notice several important features. For lattice packings, since log *δ* ≤ 0, there is no more than one center per unit sphere packing volume for *D* ∈ [0, 30]. There is exactly one in *D* = 0 and *D* = 24: these are the densest sparse packings. In higher dimensions (*D >* 30), there can be more than one center per unit sphere packing volume.

Comparing Fig 3A to Fig 3B, we encounter a puzzle. The center density in *D* = 20 is larger than the center density in *D* = 16. Why then are there more 4 base cutters than 5 base cutters? There are at least three explanations. Evolutionary selection should increase the molecular machine capacity by finding relative maxima of the sphere packing center density but evolution also minimizes the expenditure of resources by a cell. On average, an enzyme that recognizes a shorter sequence requires less protein structure and so requires less energy to synthesize. Perhaps the gain in information density at dimension 20 compared to dimension 16 is insufficient to offset the greater energy cost. In contrast, the significant improvement of information density at dimension 24 may provide a superior benefit to the organism despite the extra expenditure in energy and this may explain why there are more 6 base enzymes than 4 base enzymes. A second consideration is the number of target restriction sites needed on foreign DNA. Larger target DNAs would be best digested less frequently so that the restriction enzyme spends less time on small regions and conversely some enzymes may be targeted to smaller DNAs, leading to a smaller dimensionality. A third factor is the ease of evolving recognition patterns. Protein dimerization allows the creation of a 4 base cutting restriction site from two half sites. Five base recognition is probably more difficult to evolve since the central base has to be handled separately or the entire site has to become asymmetric.

The best center density *δ* in 16 dimensions has the same value as the best center density in 17 dimensions (Fig 3) [26]. Since higher dimensionality allows lower error rates [15], why isn’t 17 dimensions used to provide more accurate restriction? A 17 dimensional packing would take 4.25 bases and the 0.25 bases could be contacted in the center of the site. Dimeric proteins use less protein structure than a monomer and so require smaller DNA coding, but only one of the two monomers could contact the center at a time. Perhaps this awkward wasteful situation is unfavorable compared to using 16 dimensions. In fact, odd dimensions are avoided by restriction enzymes in general (Table 1). Another possibility is that the huge number of known packings in 16 dimensional space (10^7^, [59]) overwhelms a smaller number of packings in 17 dimensions. Similar considerations apply to dimensions 7 and 8, where the center densities are equal. Here, because 8 is even, the *E*_8_ lattice is known to be highly symmetric, and the error rates are smaller, the preference for *D* = 8 over *D* = 7 is clear.

According to Fig 3A there are several hundred restriction enzymes that operate in 16, 18 and 20 dimensional coding spaces. Just as the *E*_8_ and Leech lattices allow for dense sphere packings in 8 and 24 dimensions, evolution may select similarly dense packings in other even dimensions, especially those divisible by 4. There may be best packings in these dimensions that have not been discovered yet; this is an open problem in mathematics.

Evidently many restriction enzymes have discovered the Leech lattice, but does this merely reflect divergent evolution from a common ancestor? Many restriction enzymes have widely different structures [2], suggesting convergent evolution. Perhaps a deeper understanding of the coding spaces will help to classify these enzymes. In addition, the coding spaces of restriction enzymes may provide a fertile ground for precise quantitative analysis of population genetics and theoretical evolutionary biology since much is known about sphere packing in high dimensions [26]. Though we have found a strong correlation between the high dimensional sphere center packing landscape and restriction enzyme information content preferences, this still leaves the important task of understanding how the codes are implemented by the protein structures as a major problem for coding theorists and biologists.

## 11 Transcription factors use high dimensional fractal coding

Experimental evidence has been obtained indicating that for the transcription factor Fis, the specific DNA binding mode differs sharply from non-specific DNA binding since there is a break in the binding curve at zero bit binding sites [16, 92, 93]. Shannon pointed out that a mapping from a high dimensional space to lower dimensions creates discontinuities [15], so this result suggests that Fis functions in a high dimensional coding space close to 2 × 7.86 ± 0.27 = 15.72 ± 0.54 ≈16 dimensions [94].

For nucleic acid recognizing molecules that have specific sites on the genome, such as transcription factors, the information in the binding sites, *R*_sequence_, evolves to match the information needed to locate the binding sites,

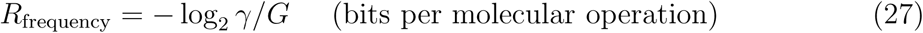

where *G* is the number of potential binding sites in the genome and *γ* is the number of specific binding sites [23, 24]. In the case of restriction enzymes equation (27) does not apply since there are no specific binding sites in the foreign DNA attacked by these enzymes and *γ* does not have a particular value. However, the principle that *R*_sequence_ ≈ *R*_frequency_ does apply to many other genetic systems such as transcription factors [23], promoters [95,96], ribosome binding sites [97,98], and mRNA splicing [99]. In general *G/γ* is not an exact power of two, so *R*_frequency_, and therefore *R*_sequence_ is usually not an integer. Equation (21) then implies that for an optimal molecular machine in which *D* = 2*R*, the dimension of the binding sites will not be an integer. Objects with non-integer dimensions are called fractals [100–102]. As shown in Table 1, many restriction enzymes may also have fractal dimensions, although more close inspection using sequencing technologies may be required to confirm the observation [103]. How a molecular coding system can have non-integer dimensions and the possible applications of such high dimensional fractal codes for communication systems remain to be investigated.

## Acknowledgments

TDS thanks N. J. A. Sloane for useful discussions about the normalization of the sphere packing density function, Rich Roberts for useful discussions about REBASE and for pointing out that Type I restriction enzymes differ from Type II, Michael Smith for suggesting that sphere packing density may be an adaptive landscape, Eckart Binde-wald, Misha Kashlev, Ryan Shultzaberger and Randall Johnson for comments on the manuscript and the Advanced Biomedical Computing Center (ABCC) for support. VJ thanks the U.S. National Cancer Institute Werner H. Kirsten Student Intern Program, Soren Brunak, and the Technical University of Denmark for hospitality during initial stages of this project in 1994.

## Data and materials availability

see Table 1 and Fig 3.

